# A conserved somatic sex determination cascade instructs trait-specific sexual dimorphism in horned dung beetles

**DOI:** 10.1101/2024.10.10.617608

**Authors:** London C. Mitchell, Armin P. Moczek, Erica M. Nadolski

## Abstract

Sex-specific trait expression represents a striking dimension of morphological variation within and across species. The mechanisms instructing sex-specific organ development have been well studied in a small number of insect model systems, suggesting striking conservation in some parts of the somatic sex determination pathway while hinting at possible evolutionary lability in others. However, further resolution of this phenomenon necessitates additional taxon sampling, particularly in groups in which sexual dimorphisms have undergone significant elaboration and diversification. Here, we functionally investigate the somatic sex determination pathway in the gazelle dung beetle *Digitonthophagus gazella*, an emerging model system in the study of the development and evolution of sexual dimorphisms. We find that RNA interference (RNAi) targeting *transformer (tra)* caused chromosomal females to develop morphological traits largely indistinguishable from those normally only observed in males, and that *tra*^RNAi^ is sufficient to induce splicing of the normally male-specific isoform of *doublesex* in chromosomal females, while leaving males unaffected. Further, *intersex*^RNAi^ was found to phenocopy previously described RNAi phenotypes of *doublesex* in female but not male beetles. These findings match predictions derived from models of the sex determination cascade as developed largely through studies in *Drosophila melanogaster*. In contrast, *transformer2*^RNAi^ resulted in larval mortality and was not sufficient to affect *doublesex* splicing, whereas RNAi targeting *Sex-lethal* and two putative orthologs of *hermaphrodite* yielded no obvious phenotypic modifications in either males or females, raising the possibility that the function of a subset of sex determination genes may be derived in select Diptera and thus non-representative of their roles in other holometabolous orders. Our results help illuminate how the differential evolutionary lability of the somatic sex determination pathway has contributed to the extraordinary morphological diversification of sex-specific trait expression found in nature.

## Introduction

Sexual dimorphism, or the presence of trait differences between the sexes, is widespread across eukaryotes and represents one of the most striking dimensions of phenotypic variation both within and across species. Sexual dimorphisms are among the fastest evolving traits and as such constitute the trait class most often used to morphologically distinguish closely related species. Furthermore, because sexual dimorphisms may involve traits greatly elaborated in or even limited to one sex only, sex-specific trait expression has been hypothesized to be an important early stepping-stone in the initiation of novelty in development and evolution. Consequently, secondary sexual traits used as weapons during mate competition and the exaggerated ornaments critical for male displays and female choice have been traditional foci of functional ecology and evolutionary theory (Andersson & Simmons 2006). However, the mechanisms underlying their development have remained generally understudied, except in select model organisms with tools for genetic manipulation. While these investigations have been critical in advancing our understanding of the developmental genetic mechanisms underlying *e.g.* somatic sex determination, they also encounter limitations, especially in insects. Developmental genetic model systems such as the fruit fly *Drosophila melanogaster* and the red flour beetle *Tribolium castaneum* possess only relatively modest degrees of morphological sexual dimorphism generally not reflective of the extraordinary degrees of elaboration and diversification we see in nature, and their ecological or other significance may be difficult to discern in the wild. Furthermore, even within the small sampling of model taxa undertaken to date, a surprising diversity of mechanisms underlying sex-specific development have been documented, suggesting that investigation of additional taxa may be warranted (Hopkins and Kopp 2021). Therefore, here we investigate the molecular genetic basis of morphological sexual dimorphisms in a non-traditional insect model system with striking and highly diversified degrees of morphological sexual dimorphisms.

The mechanisms instructing sex-specific organ formation and/or elaboration have been particularly well studied in the fruit fly *Drosophila melanogaster*. In female flies, the so-called sex determination cascade begins with factors activated by the double X chromosome dosage, which result in the splicing of the active isoform of Sex-lethal (Sxl), which in turn results in the splicing of the active isoform of Transformer (Tra, Salz & Erickson 2010). The resulting Tra protein, in combination with the constitutively spliced Transformer2 (Tra2) protein, then regulate the splicing of the female isoform of Doublesex (DsxF; Hoshijima *et al*. 1991). Along with two necessary cofactors, Intersex and Hermaphrodite, the female Doublesex protein then instructs the development of female phenotypes (Li & Baker 1998, Garret-Engele *et al*. 2002). In male flies, which possess XY sex chromosomes, Sex-lethal is instead spliced into an inactive form, which results in the baseline splicing of an inactive form of Transformer. Subsequently, only the male Doublesex isoform (DsxM) is spliced, which does not require any co-factors to regulate development of male phenotypes. In male (XY) flies, expression of *transformer2, intersex, and hermaphrodite* can be detected, but functional analyses indicate that these genes are not involved in specifying somatic sex in males (Figure 1A). Outside of flies, members of the *doublesex-mab3-related transcription factor* (DMRT) gene family have been implicated across holometabolous insect orders as a conserved genetic switch with sex-specific isoforms regulating sexual differentiation and sexually dimorphic development throughout the organism (silkworm, Ohbayashi *et al*. 2002; honeybee, Cho *et al*. 2007; jewel wasp, Beukeboom & van de Zande 2010, red flour beetle, Shukla & Palli 2012a; for evolution of this mechanism and Hemipteran data see Wexler *et al*. 2019).

**Figure 1.**
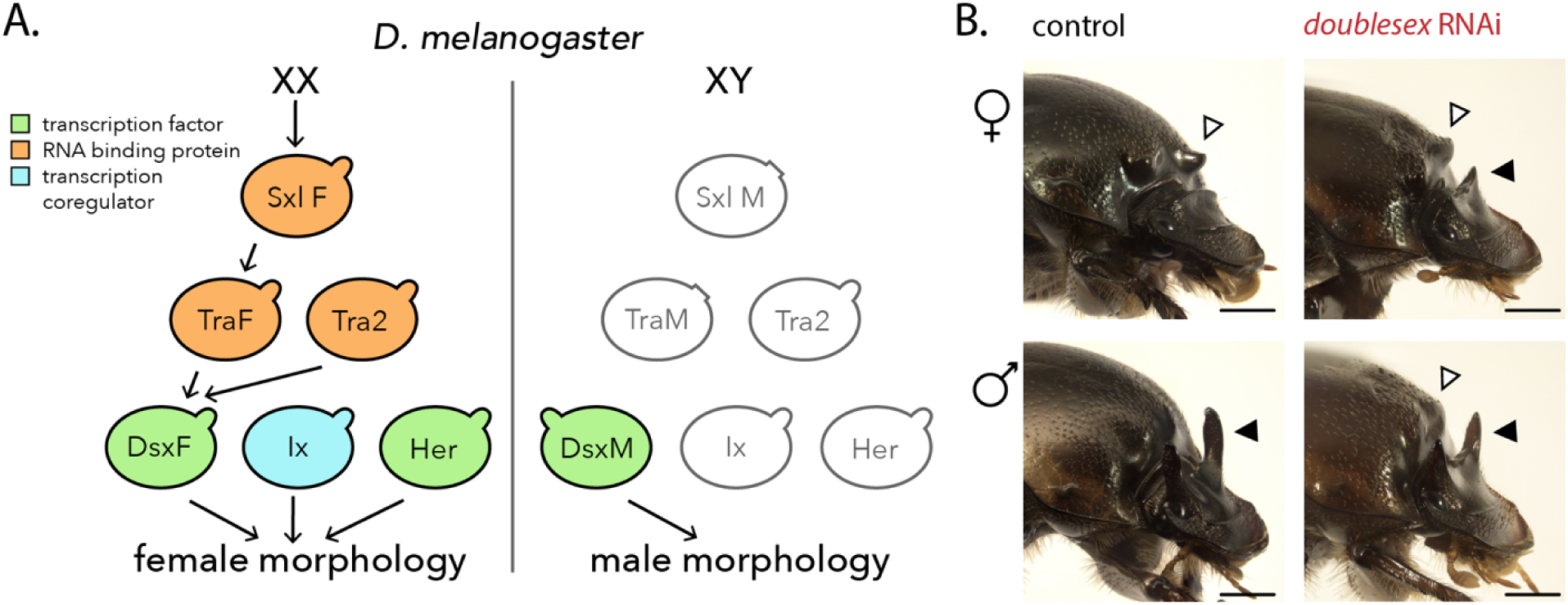
Schematic of core sex determination cascade in *Drosophila melanogaster* and the conserved role of *doublesex* in *Digitonthophagus gazella*. (A) The female sex determination cascade is depicted on the left side of the diagram: animals with two X chromosomes regulate splicing of the active isoform of Sex-lethal, which splices the active isoform of Transformer. This protein, in combination with Transformer2, regulates the splicing of the female isoform of Doublesex. Along with two required cofactors, Intersex and Hermaphrodite, the female Doublesex protein regulates development of female phenotypes. The male cascade is depicted on the right side of the diagram: in animals with XY sex chromosomes, Sex-lethal is spliced into an inactive form of the protein, which results in the splicing of only the inactive form of Transformer. In turn, this results in the transcription and splicing of only the male Doublesex isoform, which alone regulates development of male phenotypes. In XY flies, expression of transformer2, intersex, and hermaphrodite can be detected, but functional analyses indicate that these genes are not involved in the male sex determination cascade. For information on the X-linked signal elements (XSEs) that are the primary links between X chromosome dosage and regulation of Sex-lethal splicing in *Drosophila*, see Salz & Erickson 2010. For information on the downstream targets of Doublesex isoforms, see Clough et al. 2014 (*Drosophila melanogaster*) and Ledón-Rettig et al. 2017 (*Onthophagus taurus*). (B) Wildtype adult male and female *Digitonthophagus gazella* display multiple novel sexually dimorphic traits: (i) male-specific paired, straight posterior head horns (black arrowhead, bottom left), (ii) female-specific paired, rounded prothoracic protrusions (white arrowhead, top left), and (iii) differences in tibiae shape and size with females possessing short, wide forelegs woth large, wide tibial teeth and males displaying much longer, more slender tibiae with small, more rounded tibial teeth (not shown). RNAi targeting all *doublesex* isoforms eliminates these sex differences, generating beetles with intermediate phenotypes by (i) inducing horn formation in females but decreasing horn size in males, (ii) inducing prothoracic protrusions in males but decreasing their size in females, and (iii) decreasing foretibia length in males and increasing it in females (not pictured, see Rohner et al 2021). These *dsx*^RNAi^ phenotypes in D. gazella and other studies (Kijimoto et al 2012) establish conservation of its function as a key sex-determination factor in horned beetles. Scale bars = 1mm.

Despite this deep conservation of DMRT genes acting as a genetic switch via the expression of alternative, sex-specific isoforms, the pathway upstream of this switch appears to be much more evolutionarily labile. Across arthropods, significant variation exists both at the chromosomal level – with an XX-XY system in many flies and beetles, a ZW-ZZ system in butterflies, and haplodiploidy in hymenopterans – and at the level of the cascade’s first molecular signal, for example via sex-dependent activation of the *Sex-lethal (Sxl*) locus in *Drosophila* (Salz & Erickson 2010) or sex-specific expression of a piRNA, *feminizer,* in *Bombyx* (Kiuchi *et al*. 2014). Yet despite the appreciable diversity in these upstream mechanisms, in all species studied thus far the pathway then converges on the sex-specific splicing of *dsx*, which is relayed throughout the body as a signal for sex-specific differentiation and growth (Verhulst and van de Zande 2015). Likewise, *transformer*, the splicing factor directly upstream of *dsx,* has also been demonstrated to be functionally conserved across all orders studied thus far (Geuverink & Beukeboom 2014, Verhulst *et al*. 2010); however, the mechanisms by which its transcription is activated and maintained has been found to differ across groups (Gempe *et al*. 2009, Verhulst *et al*. 2010a). Thus, data to date suggest a remarkable differential evolutionary lability of the somatic sex determination cascade, characterized by striking conservation in some parts and rapid evolution in others. However, further characterization of this phenomenon and its contributions to organismal diversity will require additional taxon sampling, in particular in groups in which sexual dimorphisms have undergone significant elaboration and diversification. Here, we investigate the functional significance of an array of members of the somatic sex determination cascade in the gazelle dung beetle, an emerging model system in the study of the development and evolution of sexual dimorphisms.

Onthophagine beetles represent a powerful model system to investigate the molecular, ontogenetic, and evolutionary underpinnings of sex-specific development due to the diversity of experimentally accessible sexual dimorphisms within and among taxa (Ledon-Rettig *et al*. 2016, Davidson *et al*. 2023). Onthophagine beetles possess an XX-XY sex chromosome system, typical of most Coleoptera. Past functional genetic studies have confirmed the role of a single ortholog of *doublesex* as a sex-specifically spliced transcription factor regulating sex-biased development in the bull-headed dung beetle *Onthophagus taurus*, its close relative *O. sagittarius*, and the more distantly related *Digitonthophagus gazella* (Figure 1B, see Kijimoto *et al*. 2012, Casasa *et al*. 2020, Rohner *et al*. 2021). Additional studies have investigated downstream targets of doublesex acting as trait-specific effector genes (Ledon-Rettig *et al*. 2017). However, the role and significance - if any - of other members of the sex determination pathway in the regulation of sexual dimorphisms remain to be investigated. Here, dung beetles offer a promising opportunity to investigate the means by which information about chromosomal sex is transmitted to instruct the development of various degrees of sexual dimorphism. In this study we focused on the role of five cardinal members of the sex determination pathway as established in *D. melanogaster,* and how they may be regulating the development of sexually dimorphic traits in the gazelle dung beetle *Digitonthophagus gazella*.

Specifically, we focused our efforts on six morphological traits exhibiting varying degrees of sexual dimorphism in *D. gazella*: (i) paired posterior head horns present in males but entirely absent in females (Fig 1B); (ii) paired prothoracic protrusions present in females but absent in males (Fig 1B); (iii) foretibiae adapted for sex-specific behaviors (females possess a short, wide foreleg with large tibial teeth used for subterranean tunneling, while the male foreleg is much thinner, conspicuously elongated, and with short tibial teeth used during copulatory encounters, Fig 2B); (iv) relative length of the pygidium (the sclerite covering the opening of the genital tract) and the posterior-most abdominal sclerite (Fig S1); (v) the presence of obvious bilateral arched cuticular grooves on the inside of the pygidium in females but not males (Fig S2); and (vi) the sex-specific interior genital components: the male aedeagus and the female vagina and *receptaculum seminis* or spermatheca (Fig S3). Together, these traits comprise a spectrum of ‘degrees of sexual dimorphism’ in the level of morphological differences discernible between adult males and females. Focusing on these six traits we then sought to identify and functionally characterize the role of key sex determination pathway genes upstream of *doublesex* (*Sxl, tra1, tra2, hermaphrodite,* & *intersex)*. Specifically, we sought to establish whether cardinal sex determination pathway members characterized in other insect taxa play conserved or divergent roles in horned dung beetles by acting as regulators or cofactors of *doublesex* using a combination of functional genetic analyses and by directly investigating *doublesex* splicing in a subset of our treatments.

**Figure 2.**
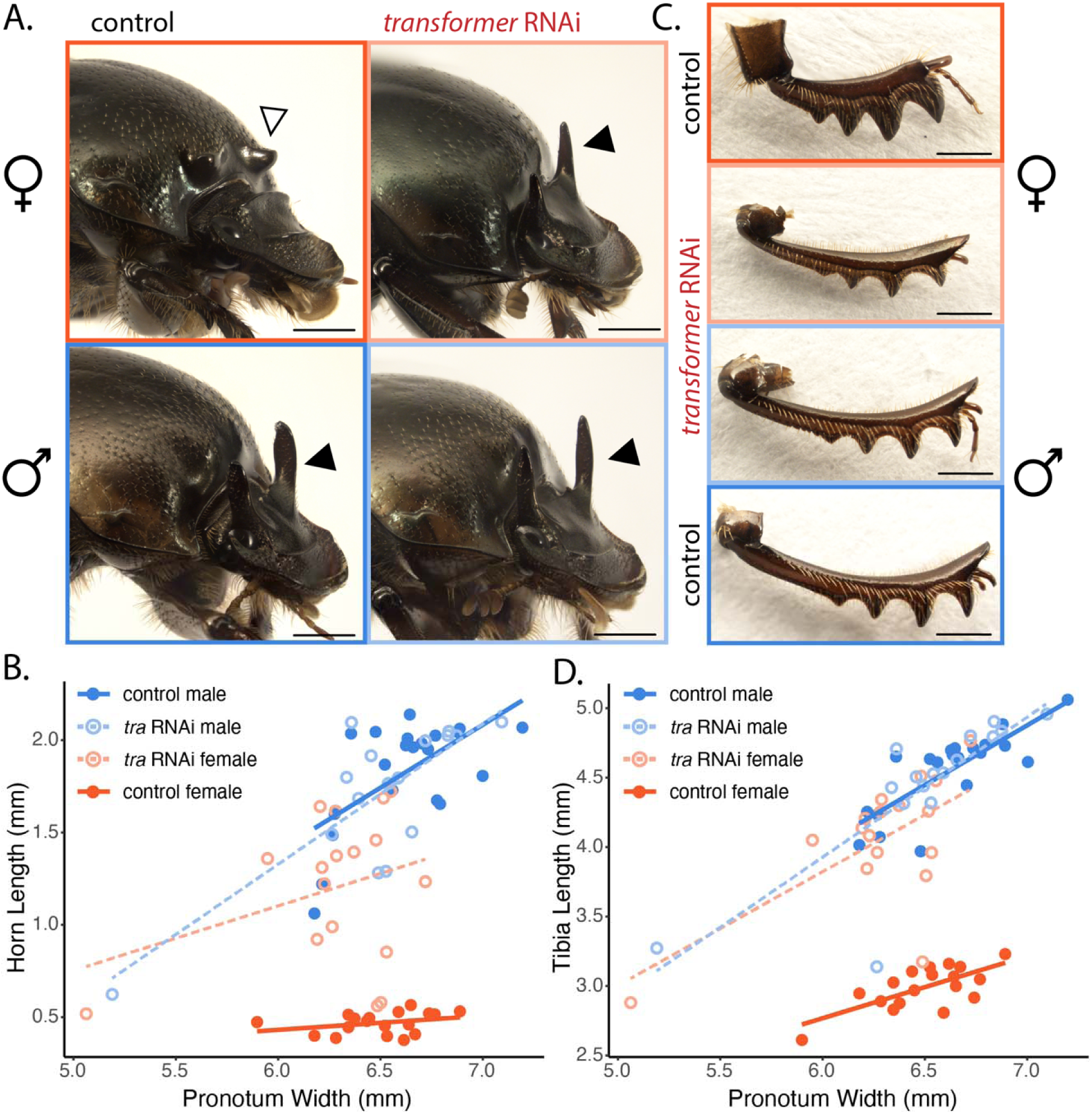
Effects of *transformer* RNAi on adult *D. gazella.* Representative animals obtained after control injections (black labels) and *tra* dsRNA injections (red labels) illustrating head horns (highlighted with black arrowheads) and prothorax protrusions (highlighted with white arrowheads) (A). Horn length for horned individuals and head ridge height for hornless individuals plotted against pronotum width, a proxy measurement for overall body size, for all individuals (B). Results indicate that *Dg-tra*^RNAi^ did not affect adult male traits, but in females substantially reduced prothoracic protrusions and induced conspicuous, paired ectopic head horns (panel A upper right; black arrowhead, panel B light red data points). Also shown are representative male and female fore tibiae after control and *tra* dsRNA injections (C) and corresponding measurements of tibial length (D). Both length and tooth size of female foretibiae transformed to resemble the longer, thinner morphology normally only observed in males, while male *Dg-tra*^RNAi^ individuals were unaffected. Scale bars = 1mm.

## Methods

### Beetle husbandry

*Digitonthophagus gazella* individuals were collected near Barber County, Kansas and reared in a lab colony as described previously (Moczek & Nagy 2005). Reproductively active adults were transferred from the colony into a breeding container and allowed to reproduce. Eclosed larvae were transferred from their natal brood balls into twelve-well plates and provided with organic cow dung from Marble Hill Farm (Bloomington, IN) as described in Shafiei *et al*. (2001) and kept in 16:8h light/dark cycle at 28°C pre- and post-injection treatments until eclosion. Eclosed adults were sacrificed and preserved in 70% ethanol.

### BLAST ortholog identification

We generated a custom BLASTP database for the *D. gazella* proteome (Davidson & Moczek 2024) using BLAST (Altschul *et al*. 1990). Query protein sequences of known *D. melanogaster* sex determination genes were collected from Flybase (version FB2024_02, Öztürk-Çolak *et al*. 2024), and their predicted orthologs from the *Tribolium castaneum* and *O. taurus* genomes were collected from OrthoDB (Kuznetsov *et al*. 2023). The query sequences were used to search the custom *D. gazella* proteome database to identify target protein sequences for Sex-lethal, Transformer, Transformer2, Intersex, and Hermaphrodite. We established an e-value cutoff of 1×10^-5^ for selecting the best hit from each list of potential targets, and prioritized targets that appeared as the top hit above the e-value cutoff for all queries.

### Double-stranded RNA synthesis for RNA interference

Gene fragment constructs (Table S1) and fragment-specific primers (Table S2) were designed using the *D. gazella* reference genome and ordered from Integrated DNA Technologies, Inc. Fragments of each gene suitable as targets for RNA interference were chosen by using BLAST to query 250bp portions of each gene against a custom *D. gazella* transcriptome database and selecting those with zero off-target hits. Note that *transformer* and *transformer2* are not paralogs despite the naming convention. Synthesis of double-stranded RNA (dsRNA) for gene knockdown via RNA interference was performed using a protocol optimized for coleopteran larvae (Philip & Tomoyasu 2011). In brief, for each gene, the DNA template for dsRNA synthesis was synthesized via PCR to add T7 RNA polymerase binding sequences flanking the gene fragment. These constructs were purified using a Qiagen QIAquick PCR Purification kit. In vitro transcription of dsRNA was performed using an Ambion MEGAscript T7 kit, and each target-specific dsRNA product was purified using an Ambion MEGAclear kit and an ethanol precipitation step.

### Injection of constructs for RNA interference

dsRNA constructs were diluted to 1 ug/uL with injection buffer (Philip & Tomoyasu 2011). Prior to injection, we sexed each larva to confirm chromosomal sex as described in Moczek and Nijhout (2002). During the late second to third larval instar stage, chromosomal male larvae exhibit increasingly prominent genital primordia visible underneath the cuticle in the ventrocaudal abdomen. These tissues are absent in chromosomal females, allowing for unambiguous sexing of each larva (Fig S4). A 3µl dose of dsRNA targeting a single gene was injected through the abdominal cuticle into the hemolymph of sexed second and third-instar larvae using a Hamilton brand syringe and small 32-gauge removable needle. Control individuals were randomly selected from each round of developing larvae and injected with pure injection buffer. Previous work in this system has shown that injection of either pure buffer or nonsense RNA can serve as suitable controls for dsRNA injection, as neither buffer-nor nonsense RNA-injected adults show any detectable phenotypic differences compared to wildtype adults (Moczek & Rose 2009, Kijimoto *et al*. 2012).

### Phenotype scoring and photography

We analyzed the phenotypes of RNAi and control individuals post eclosion. We specifically focused our analysis on the following six sexually dimorphic traits in control and RNAi individuals. First, we assessed (i) the presence or absence of head horns (normally only seen in wildtype males), (ii) prothoracic protrusions (normally only seen in wildtype females), and (iii) the shape and size of foretibiae (drastically elongated and thinner in males compared to females). In addition, we assessed the morphology of the pygidial flap, a sexually dimorphic projection attached to the terminal abdominal segment which covers access to the genital track. Specifically, we evaluated the (iv) exterior of the pygidium and the closure it forms against the subsequent abdominal sclerite, which is consistently sexually dimorphic across the Onthophagine clade: males possess an elongated pygidium, resulting in a conspicuous medial narrowing of the closure between pygidium and the neighboring abdominal sclerite, whereas females display a consistent spacing between pygidium and neighboring sclerite. Additionally, we evaluated the presence of (v) bilateral arched cuticular grooves on the interior of the pygidium, which are present in females and nearly absent in males. Components of the (iv) internal genitalia were also dissected from representative individuals of both sexes for each sample group, following the terminology of Roggero *et al*. 2017; specifically, males were examined for the presence of an aedeagus comprised of a proximal phallobase and distal parameres ending in conspicuous paired projections, whereas females were examined for the presence of a vagina and spermatheca. Representative individuals from each sample group were photographed using a Leica MZ16 microscope with a PLANAPO 2.0x objective (Bannockburn, IL, USA) and a PixeLINK PL-D7912CU-T digital camera (Scion, Frederick, MD, USA); multiple photos of each sample were taken across different planes of focus and overlaid using Adobe Photoshop.

For a subset of RNAi manipulations and body regions, allometric measurements were taken to enable a quantitative assessment of the RNAi phenotypes observed. For these sample groups, every RNAi and control individual was photographed as described above. For control and RNAi individuals with head horns, horn length was measured by taking one linear measurement from the posterior corner of the left eye to the tip of the left horn (as in Rohner *et al*. 2023). For hornless individuals, head ridge height was measured using one linear measurement from the corner of the eye to the left vertex of the head ridge. Tibia length was measured using one linear measurement from the farthest point on the convex edge of the proximal tibial joint with the femur to its distal tip next to the tarsal segments (as in Rohner *et al*. 2023). Finally, we used pronotum width as a proxy for overall body size in this species which was measured using one linear measurement at the widest point across the pronotum (as in Rohner *et al*. 2023). Trait lengths were measured using Fiji (Schindelin *et al*. 2012) and plotted using the R package ggplot2 (V3.5.1, Wickham 2016).

### Doublesex RT-PCR following tra1 & tra2 manipulation

To assess if *Dg-tra1*^RNAi^ and *Dg-tra2*^RNAi^ are sufficient to affect *doublesex* isoform splicing in *D. gazella*, additional *Dg-tra1*^RNAi^, *Dg-tra2*^RNAi^, and control injected larvae were reared for RNA extraction and RT-PCR targeting the *doublesex* coding region. Larvae were injected as described above, and upon metamorphosis the head and thoracic region was dissected and stored in TriZOL at –20°C. RNA extraction was performed using a Zymo Direct-zol RNA Miniprep kit. Extracted RNA samples were used (i) to perform RT-PCR with primers targeting the *doublesex* coding region (Table S2) using a ThermoFisher SuperScript TM IV One-Step RT-PCR kit to assess isoform size, and (ii) sent for Sanger and/or Illumina amplicon sequencing based on the number of bands per sample, to assess isoform sequence. Illumina amplicon sequencing was performed after Nextera Small Genome DNA Library preparation on a NextSeq 2000 with a P2 2×100 flow cell. Sequencing data were pre-processed using FastQC (Andrews 2010) and Trimmomatic (Bolger *et al*. 2014), then using Trinity (Haas *et al*. 2013) to perform genome-guided de-novo transcriptome assembly to assemble transcripts from sequencing reads. The assembled sequences were then aligned to *O. taurus* and *O. sagittarius* doublesex isoform sequences from Kijimoto *et al*. (2012) and manually annotated to determine predicted exon boundaries.

## Results

We sought to investigate the function of the cardinal sex determination genes *Sex-lethal, transformer, transformer2, intersex,* and *hermaphrodite* in the sexually dimorphic gazelle dung beetle, *D. gazella,* through identification of target homologs, assessment of RNAi phenotypes resulting from RNA interference targeting each gene, and examination of doublesex splicing patterns following a subset of RNAi treatments. Below we discuss each of our findings in turn.

### BLAST Ortholog Identification

We identified *D. gazella* target homologs for all five genes of interest using BLAST with queries from *D. melanogaster, T. castaneum,* and *O. taurus.* For the Intersex, Sex-lethal, and Transformer2 proteins, there was an unambiguous top hit matching to each query (Table S2). For the Transformer protein, the top hits diverged across the queries, so an additional query from *Trypoxylus dichotomus* (a closer relative than *D. melanogaster* or *T. castaneum*) was used, and the top hit found to be matching both the *T. dichotomus* and *O. taurus* query was used as the target gene (Table S2). For Hermaphrodite, which has not been experimentally validated outside of the *Drosophila* genus, predicted homology of the hits to each query was far lower. We therefore carefully examined those sequence hits appearing above our e-value cutoff for both *T. castaneum* and *D. melanogaster* and chose two sequences with the highest conservation as target genes to proceed with functional genetic characterization (Table S2), as detailed next for each target gene.

### Sex-lethal RNAi

RNA interference targeting *Dg-Sxl* resulted in no change in phenotype in any of the sexually dimorphic body regions, nor any obvious morphological defects in monomorphic body regions. Specifically, the head horn, prothorax, foreleg, and genital phenotypes of *Dg-Sxl*^RNAi^ injected individuals matched those of control individuals in both sexes. No other obvious morphological defects were observed (Fig S5A, Table S3).

### Transformer RNAi

*Dg-tra* RNA injection resulted in a dramatic degree of masculinization of the female head, prothorax, and foreleg. Specifically, females injected with the *Dg-tra*^RNAi^ construct developed prominent head horns similar to those of same-sized males (Fig 2A-B), featured a smooth prothorax missing bilateral protrusions otherwise typical of female morphology (Fig 2A), and developed skinnier, longer, male-like fore tibiae (Fig 2C-D, Table S3). Additionally, the shape of the pygidium closure – i.e. the abdominal sclerite covering the entrance to the genital tract – and the internal pygidial cuticle morphology underwent significant masculinization in *Dg-tra*^RNAi^ females; specifically, the closure of the pygidium of RNAi females narrowed significantly in the medial abdomen, matching control males (Fig S1B), and the internal pygidial cuticle grooves were conspicuously smaller than those of control females (Fig S2B). Finally, typical female internal genitalia were absent, but in contrast to the other body regions, masculinization in this body region was not observed – i.e. no male-like internal genital features were found (Fig S3). In stark contrast, RNAi targeting *Dg-tra* in male larvae did not result in any changes of the adult phenotypes of the head, prothorax, foretibiae (Fig 2), closure of the pygidium or the internal pygidial cuticle (Fig S1B, S2B); similarly, the internal genitalia of *Dg-tra*^RNAi^ males matched those of control males without exception (Fig S3).

### Transformer2 RNAi

RNAi targeting *Dg-tra2* caused 93% lethality throughout the larval and pupal stages (Table S3). Surviving adults showed no obvious phenotypic differences compared to control-injected or wildtype individuals, in either the sexually dimorphic regions of interest or otherwise (head, prothorax, foretibiae: Fig S5B, pygidium closure: Fig S1E, internal pygidium morphology: Fig S2E). Note that the single surviving *Dg-tra2*^RNAi^ male was small-bodied, and could not be size matched to other control and RNAi males; *D. gazella* male head horns and fore tibiae scale with body size in a hyper-allometric manner, so the relatively small horns and short tibiae of this *Dg-tra2*^RNAi^ male can most likely be accounted for solely by body size differences and therefore do not represent an RNAi phenotype.

### Intersex RNAi

Female individuals injected with the *Dg-ix*^RNAi^ construct exhibited a pair of small but obvious head horns (Fig 3A-B), reduced prothoracic protrusions (Fig 3A), and moderately longer, thinner forelegs compared to control-injected females (Fig 3C-D, Table S3). In the posterior abdomen, the pygidium and inner pygidial grooves displayed a slight phenotypic shift toward a more masculine shape (Fig S1C, Fig S2C). Male *Dg-ix*^RNAi^ individuals matched controls across all phenotypes: head, prothorax, foretibiae (Fig 3), pygidium closure (Fig S1C), internal pygidium morphology (Fig S2C), and internal genitalia (Fig S3).

**Figure 3.**
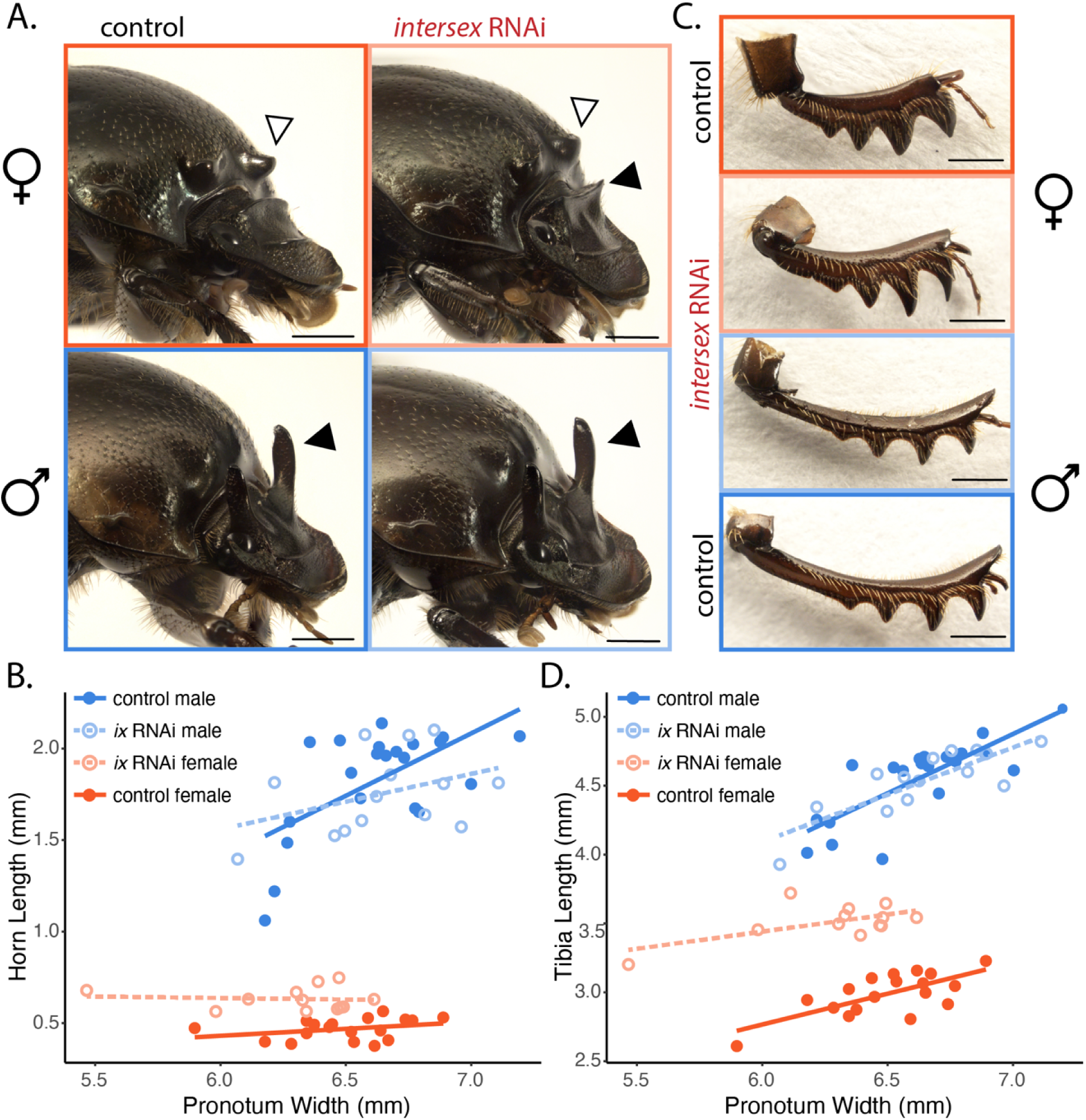
Effects of *intersex* RNAi on adult *D. gazella.* Representative animals obtained after control injections (black labels) and *ix* dsRNA injections (red labels) illustrating head (horns highlighted with black arrowheads) and prothorax (protrusions highlighted with white arrowheads) (A). Horn length for horned individuals and head ridge height for hornless individuals plotted against pronotum width, a proxy measurement for overall body size, for all individuals (B). Results indicate that *Dg-ix*^RNAi^ did not affect adult male traits, but in females the treatment moderately reduced prothoracic protrusions, and induced small ectopic head horns (panel A upper right; black arrowhead, panel B light red data points). Also shown are representative male and female fore tibiae after control and *ix* dsRNA injections (C) and corresponding measurements of tibial length (D). *Dg-ix*^RNAi^ modestly masculinized the foretibiae by generating slightly elongated, thinner morphology, intermediate between control males and females, while male *Dg-tra*^RNAi^ individuals were unaffected. Scale bars = 1 mm.

### Hermaphrodite RNAi

The first potential *hermaphrodite* ortholog tested, *Dg-jg1708*, caused 100% lethality in the larval stage in both sexes (Table S3). In contrast, the second potential *hermaphrodite* ortholog tested, *Dg-jg4744*, resulted in only 15% lethality (Table S3). However, the surviving adults displayed no obvious phenotypic changes in the sexually dimorphic body regions or in other monomorphic traits in males or females (pygidium closure: Fig S1F, internal pygidium morphology: Fig S2F, head, prothorax, foretibiae: Fig S5C).

### Regulation of doublesex splicing

To further characterize the functions of *Dg-tra1* and *Dg-tra2* in the dung beetle sex determination pathway, we performed an RT-PCR experiment to assess if *Dg-tra1*^RNAi^ or *Dg-tra2*^RNAi^ could individually affect *doublesex* isoform splicing. After injection of *Dg-tra1*^RNAi^ and *Dg-tra2*^RNAi^ constructs, RNAi and control larvae were monitored until metamorphosis, and were then utilized for RNA extraction and RT-PCR targeting all *doublesex* isoforms. Matching results from earlier studies on the closely related *O. taurus* (Kijimoto *et al*. 2012), we found that control males expressed a single, smaller *dsx* isoform, while females express multiple larger isoforms (Fig 4, Table S4). *Dg-tra1*^RNAi^ females, in contrast, were found to lack the larger female isoforms and instead, to express the smaller male isoform, whereas *Dg-tra1*^RNAi^ males showed the same band as control males. In contrast to our results for *tra1*, RNAi targeting *tra2* did not appear to affect *dsx* splicing in either females or males.

**Figure 4.**
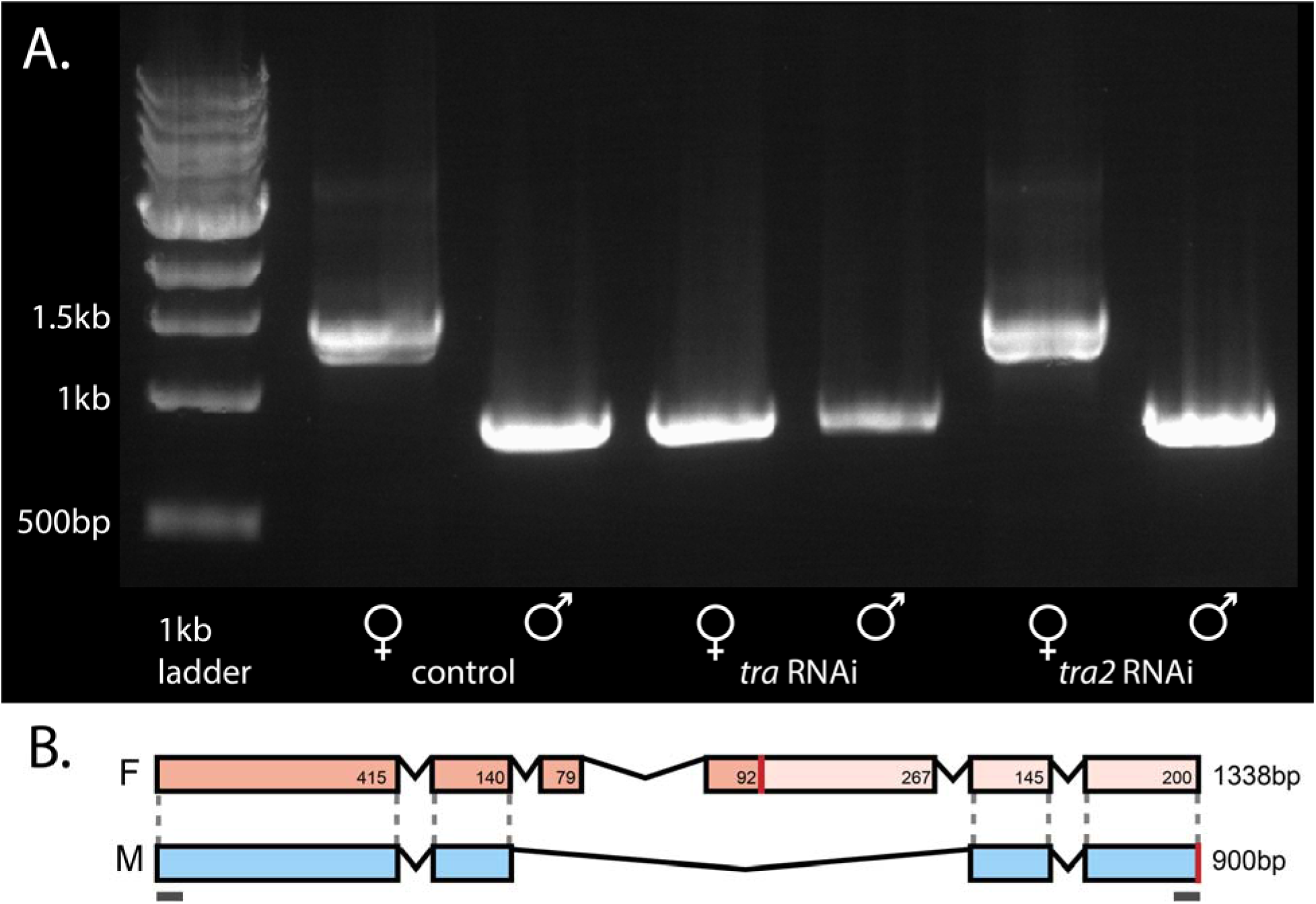
Expression and inferred structure of *D. gazella doublesex* isoforms. **(A)** RT-PCR results from left to right: 1kb ladder, control-injected female bands ∼1300-1350 basepairs, control-injected male band of ∼900bp, *Dg-tra*^RNAi^ female band of 900bp, *Dg-tra*^RNAi^ male band of 900bp, *Dg-tra2*^RNAi^ female bands of ∼1300-1350bp, *Dg-tra2*^RNAi^ male band of ∼900bp. These results confirm (i) sex-specific splicing pattern of *D. gazella doublesex* in control males and females, and (ii) the role of Transformer in simultaneously promoting the splicing of female – while preventing the splicing of the male isoform – of *dsx: Dg-tra*^RNAi^ females produced the male isoform band while female isoform bands are absent. RT-PCR primer pairs correspond to those shown in B. **(B)** Diagram of *doublesex* isoform structures in *D. gazella:* the single shorter male isoform (M) and the longer female isoform (F) sequences obtained from sequenced cDNA are indicated by rectangles, with inferred ORFs shaded either blue (male) or pink (female). Stop codons are indicated by red lines, with putative 3’ untranslated regions shown in light pink (after Kijimoto et al 2012). The horizontal grey bars indicate regions used as primers for RT-PCR.

## Discussion

In this study we sought to investigate the potential conservation of cardinal insect sex determination factors in regulating sexual differentiation in the sexually dimorphic horned beetle *D. gazella*. We performed single-gene RNAi treatments targeting male and female larvae to determine phenotypic effects in the adults and followed with RT-PCR experiments for a subset of genes to more directly assess regulatory links between *doublesex* and its potential direct upstream regulators. Four salient results emerged.

### Transformer RNAi masculinizes chromosomal females

RNA interference targeting *Dg-tra* generated females that appeared morphologically nearly indistinguishable from their chromosomal male counterparts. Head horn lengths of *Dg-tra*^RNAi^ females were in a similar size range of control and RNAi males (Fig 2A-B). Similarly, the prominent prothoracic protrusions normally observed in wildtype or control females were virtually eliminated in *Dg-tra*^RNAi^ females, leading to a simpler, rounded prothorax shape matching that of typical male *D. gazella*. Likewise, foretibiae size and shape of *Dg-tra*^RNAi^ females also transformed to the longer, thinner form normally observed only in males (Fig 2C-D). Additionally, the length between the closure of the pygidium and the adjacent abdominal sclerite decreased significantly (FigS1), and the inner pygidial grooves normally present in females decreased substantially in RNAi females, matching control male phenotypes (FigS2). These results indicate that *Dg-tra* is necessary for the regulation of female development, in line with earlier findings in *D. melanogaster* as well as more recent work in the red flour beetle *T. castaneum* (Shukla & Palli 2012) and the stag beetle *Dorcus rectus* (Gotoh *et al*. 2024). More generally, our results confirm that *Dg-tra* plays a conserved role in the sex determination pathway in horned beetles.

In partial contrast, internal female genitalia did not undergo complete masculinization after *Dg-tra*^RNAi^. While the normal female genital structures (vagina and spermatheca) were absent from *Dg-tra*^RNAi^ females as expected, no instances of ectopic male genitalia were observed. We hypothesize that this may be due to major differences in the timing of growth and differentiation of genitalia compared to our other focal traits. Specifically, the cells and tissues that give rise to the adult head including horns, prothorax, foretibiae, and abdominal sclerites do not initiate sexually dimorphic growth until the end of larval development following epidermal apolysis from the larval cuticle and entry into the larval-to-pupal molting cycle (Moczek and Nagy 2005). In contrast, the cells forming male and female internal genitalia begin proliferation and morphogenetic arrangements as early as during the transition from the second to third larval instar (SLJvácha 1992; Moczek & Nijhout 2000). Thus, the temporal window during which the signals necessary to alter sex-specific internal genital formation may have been already closed prior to our mid-third larval instar RNAi treatment, leading to only an incomplete masculinization of female genital primordia. This early specification of genital primordia relative to other appendages appears to be widespread (stag beetles: Gotoh *et al*. 2024) and may also explain the absence of RNAi phenotypes in other studies of sex determination.

### Intersex RNAi phenocopies female but not male doublesex RNAi

Our intersex RNAi treatment reduced sex differences in sexually dimorphic regions in females, but not males. Female *Dg-ix*^RNAi^ individuals exhibited a pair of small ectopic head horns, reduced but still discernible prothoracic protrusions, and slightly masculinized forelegs (Fig 3). Additionally, the distance between the closure of the pygidium and the adjacent abdominal sclerite decreased (Fig S1), and the inner pygidial grooves diminished in size in *Dg-ix*^RNAi^ females (Fig S2), all indicating a morphological state *intermediate* between typical female and male phenotypes. Collectively, Dg-ix^RNAi^ treatment thus phenocopies the effects of *Dg-doublesex*^RNAi^ in females, but not in males (Fig 1B; also see Rohner *et al*. 2021), indicating that Intersex, along with Transformer, is necessary for the proper regulation of female but not male somatic sex determination and sex-biased morphology. These findings also match the results of functional analyses in the stag beetle *Cyclommatus metallifer* (Gotoh *et al*. 2016) and are concordant with the logic of the sex determination pathway (Fig1A), which posits that the female Doublesex protein requires Intersex as a cofactor, whereas the male Doublesex protein can function alone.

### Evolutionary lability of the sex determination pathway in holometabolous insects

RNAi targeting the other putative sex determination genes examined in this study – *Dg-Sxl, Dg-tra2, Dg-jg1708*, and *Dg-jg4474* – yielded no obvious phenotypic modifications in either males or females, and regardless of whether sexually dimorphic or monomorphic body regions were examined. These null results have a range of implications.

First, the absence of *Dg-Sxl*^RNAi^ phenotypes matches results obtained in other taxa outside of Drosophilidae, including other Dipterans (*Musca domestica*, Meise *et al*. 1998; *Cyclommatus metallifer*, Gotoh *et al*. 2016; *Bombyx mori*, Niimi *et al*. 2006). This supports the possibility that despite strong conservation of the Sxl coding sequence across Holometabola, its function in Drosophilid sex determination is likely derived and divergent from that executed in other Holometabola (Sanchez 2008). Future work remains to be done to assess what other developmental roles Sxl may be playing in other taxa.

In partial contrast, previous work outside of flies has indicated some divergence in function of the Transformer2 protein across insects. While in the fruit fly it functions primarily alongside Tra1 to facilitate splicing of female *doublesex*, in other Dipterans it also has been shown to function in splicing *tra1* RNA (*Musca domestica*, Burghardt *et al*. 2005; *Lucilia cuprina*, Concha & Scott 2009; *Ceratitis capitata*, Salvemini *et al*. 2009; *Anastrepha suspensa*, Sarno *et al*. 2010). In the honeybee *Apis mellifera*, Tra2 has also been shown to splice *doublesex* in addition to the honeybee ortholog of *transformer*, *feminizer.* However, in the honeybee Tra2 was additionally documented to affect embryogenesis and resulted in lower embryonic viability (Nissen *et al*. 2012). In the red flour beetle *T. castaneum* (Tenebrionidae), *tra2* RNAi results in improper *doublesex* splicing in females, improper *tra* splicing in both sexes, and embryonic and larval lethality in both sexes (Shukla & Palli 2013). In the stag beetle *C. metallifer* (Lucanidae), *tra2* RNAi manipulations in the prepupal stage led to 100% mortality as well, and the role of this gene in sexual differentiation thus remains to be assessed (Gotoh *et al*. 2016). More generally, these data suggest that in both Hymenoptera and Coleoptera Tra2 may be executing critical functions in juvenile development independent of sex determination alone.

In this study we observed high levels of larval and pupal lethality following *Dg-tra2*^RNAi^ in *D. gazella* (Scarabaeidae), matching the lethality data obtained in *T. castaneum* and *C. metallifer*. However, no phenotypic effects were observed in the few tra2 RNAi adults that did survive to eclosion. This raises the possibility that a putative juvenile viability function of Tra2 may be ancestral in the Coleoptera, while the ancestral function in sex determination may have been lost in the family Scarabaeidae. However, the data presented here cannot exclude the alternative possibility that the lack of phenotypic effects in *tra2* RNAi adults resulted from a failure of the RNAi treatment rather than reflecting the biological function of the gene. Future work targeting the role of Tra2 across a larger sampling of beetle families could resolve these complexities, in addition to careful consideration of developmental timing during treatments in order to target appropriate developmental windows.

Lastly, our investigation into potential homologs of the *Drosophila* sex determination gene Hermaphrodite failed to yield conclusive results. Hermaphrodite homologs (with either conserved sequence or function) have yet to be identified outside the *Drosophila* genus, although the *D. melanogaster* protein is part of the large family of C2H2 zinc-finger transcription factors. BLAST ortholog identification did not yield an obvious homolog for *Dmel-her*, so we chose to proceed with functional analysis of two *D. gazella* zinc-finger transcription factors, *Dg-jg1708* and *Dg-jg4474*. *Dg-jg1708*^RNAi^ resulted in 100% mortality during the larval stage, indicating a crucial role in juvenile development, but likely also indicating an absence of true homology to *Dmel-her*. In contrast, *Dg-jg1708*^RNAi^ individuals survived to adulthood, but exhibited no obvious phenotypic effects, indicating again an absence of homology to *Dmel-her*. Currently available data suggest that *hermaphrodite* may indeed be a Drosophilid-specific gene, and that the female *Doublesex* isoform in horned beetles may only require *Intersex* as a cofactor, rather than multiple interacting cofactor proteins. Future work tracing the evolution of *Dmel-her* across Drosophilids and the entire Dipteran order could elucidate when the necessity of Hermaphrodite for female sex determination evolved.

### Transformer-Doublesex splicing mechanism is conserved

Based on the work of Kijimoto *et al*. (2012), we predicted that control males would express a single *dsx* isoform around 900 base pairs in length, while females would express multiple larger isoforms ranging from 1300-1500 base pairs. Furthermore, based on the RNAi phenotypes observed, we hypothesized that *tra1* RNAi would eliminate proper isoform splicing in females but not males.

Our RT-PCR results confirmed that doublesex is spliced in a sex-specific manner in *D. gazella*, with males expressing a single shorter isoform, and females expressing multiple longer isoforms (Fig 4). Sequencing results indicated the male isoform to be 900 basepairs in length, and the longest female isoform is predicted to be 1338 basepairs long. However, our sequencing approach was not able to resolve the full sequence of the smaller female isoform that appeared on the gel (Figure 4, Table S4). Concordant with our morphological findings, we found that *Dg-tra1*^RNAi^ females did not express bands typical of wildtype chromosomal females and instead produced the single smaller isoform characteristic of chromosomal males, suggesting that the widespread mechanism of female Doublesex splicing via Transformer is conserved in *D. gazella*. As predicted, this effect was restricted to females; *Dg-tra*^RNAi^ treatment did not affect isoform production in males (Fig 4). In contrast, *Dg-tra2*^RNAi^ did not affect splicing in either males or females, with each sex producing bands identical to their controls, in contrast to earlier findings in *Tribolium* beetles (Fig4).

These findings further illuminate why and how the reduction of sex differences through the extreme masculizination of females after *Dg-tra1^RNAi^* differs from the reduction of sex differences following *doublesex ^RNAi^*, which instead results in the production of morphologies *intermediate* to typical male and female phenotypes in both sexes: RNAi targeting *dsx* eliminates active isoforms in both sexes, thereby eliminating the sex-specific instructions for the regulation of growth and differentiation relevant for *both* male and female morphologies. In contrast, RNAi targeting *tra1* eliminates a function necessary only for the regulatory cascade underlying female-specific development, effectively re-routing the somatic sex differentiation pathway down the alternative path of masculinization via the male *dsx* isoform.

Taken together, the RNAi phenotypes and the results of the RT-PCR experiment indicate that male-type is the default splicing pattern of *dsx* in horned beetles, as the male isoform is shown here to be produced in the absence of any regulatory inputs from known members of the sex determination cascade. Earlier work in *Tribolium* has posited the existence of a dominant male “M factor” present on the Y chromosome in beetles that may act to suppress female sex determination factors to allow male development, but data confirming the existence of such a factor and how it may regulate or splice male *doublesex* have not yet materialized (Shukla & Palli 2014). Future work may elucidate whether male *doublesex* is expressed and spliced in a similar manner to ‘housekeeping’ genes. Such a mechanism may be explained by the putative evolution of the sex-specific splicing of *doublesex* in insects: in non-insect arthropods, *doublesex* produces only a single isoform that primarily regulates masculinization of male tissues through both the upregulation of masculinizing genes and the repression of feminizing genes (Kato *et al*. 2011). Current research across arthropods suggests that the derived insect mechanism may have evolved via a subdivision of ancestral functions (Kopp 2012); thus, it may be possible that the regulation of the male isoform in insects is still achieved through a more ancestral, non-sex-specific mechanism.

### Conclusion

Here, we investigated the conservation of function of insect sex determination genes in the sexually dimorphic horned beetle *D. gazella,* through single-gene RNAi experiments and an RT-PCR experiment to assess evolutionary lability in the insect sex determination cascade. Our results document that Transformer acts as a direct splicing regulator of the female isoforms of *doublesex* in horned beetles (Fig 5). Additionally, gene knockdown experiments confirmed that Intersex is required for proper female development in horned beetles, and that the role of Sex-lethal in sex determination is likely derived in Drosophilidae. Finally, this work also suggests that the role of Transformer2 in sex determination may be particularly evolutionarily labile, as it appears to have evolved a function in promoting juvenile viability in some holometabolans (Coleoptera and Hymenoptera) but not others (Diptera), and may have lost its role in sexual differentiation in Scarabaeidae after divergence from Tenebrionidae (Fig 5). One standing question that remains regards the factors upstream of Transformer in beetles; the regulatory cascade instructing Tra splicing is known in many Diptera and Hymenoptera but remains to be discovered in Coleoptera.

**Figure 5.**
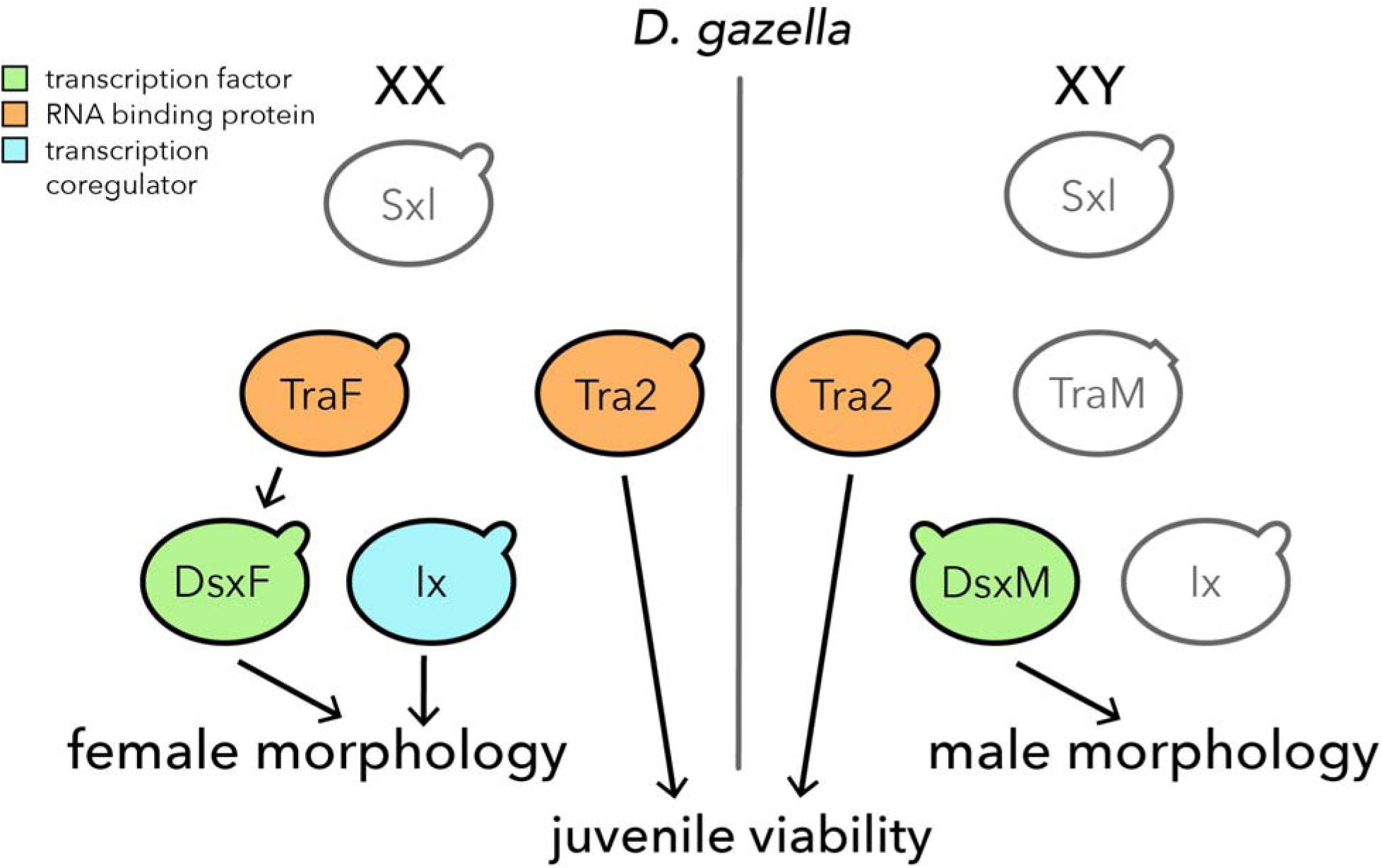
Model of the sex determination cascade in *D. gazella.* This study and past work (Rohner *et al*. 2021) have established partial conservation and partial divergence of the sex determination cascade in horned beetles compared to other holometabolous insects. The female sex determination cascade is depicted on the left side of the diagram, and the male cascade is depicted on the right. Results to date indicate that *Sex-lethal* is conserved in the beetle genome but does not function in sex determination. In female beetles, Transformer acts as a direct splicing regulator of female *doublesex* isoforms, and DsxF requires Intersex as a cofactor to regulate proper female development. Transformer2 was found to be necessary for survival through the larval and pupal stages in both males and females but did not affect *doublesex* splicing. Finally, a Hermaphrodite ortholog was not found in horned beetles. In male beetles, the male doublesex isoform can regulate male development without any known cofactors. At present, the regulatory factors upstream of Transformer in beetles are unknown.

Taken together, this work provides evidence that splicing of the male *dsx* isoform is the default state during beetle ontogeny in absence of other regulatory inputs, a finding that matches results across Holometabola. An exciting open avenue of inquiry concerns the mechanisms that regulate this default transcription of the male *doublesex* isoform. In general, the lability of both sex determination cascades on a molecular level and of sexually dimorphic traits on a morphological level suggests that greater sampling of these phenomena across orders, and of their potential regulatory links, may be poised to uncover potential important causal connections between the two. Lastly, this work contributes to a growing number of studies hinting that redeployment of deeply conserved regulatory pathways in a modular manner may be one mechanism by which even rapidly evolving traits such as those exhibiting sexual dimorphism can diversify independently in development and evolution. Future work in this exciting open area may uncover the mechanisms involved in re-deployment of the sex determination cascade in new cell types or at new developmental timepoints.

## Acknowledgements

We would like to thank Max Proctor for beetle collecting, Cale Whitworth for sharing his expertise on the *Drosophila* sex determination pathway, Anna Macagno for her advice on genitalia dissections, and the Indiana University Center for Genomics and Bioinformatics for library preparation and sequencing. Earlier drafts of this manuscript benefited from the comments of P. Davidson, R. Westwick, K. Givens, J. Jones, I. Manley, E. Pieri, and S. Kidd. Comments by two anonymous reviewers and the editor greatly improved the initial submission. This work was supported in part through generous funding from the National Science Foundation [Grant no. 2243725 and 1901680 to APM] and was performed while EMN was funded by the National Institutes of Health [T32-HD049336]. Additional support was provided by the Bloomington High School South Senior Internship Program to LCM.

