## Supplemental-figures-tables for "A conserved somatic sex determination cascade instructs trait-specific sexual dimorphism in horned dung beetles"

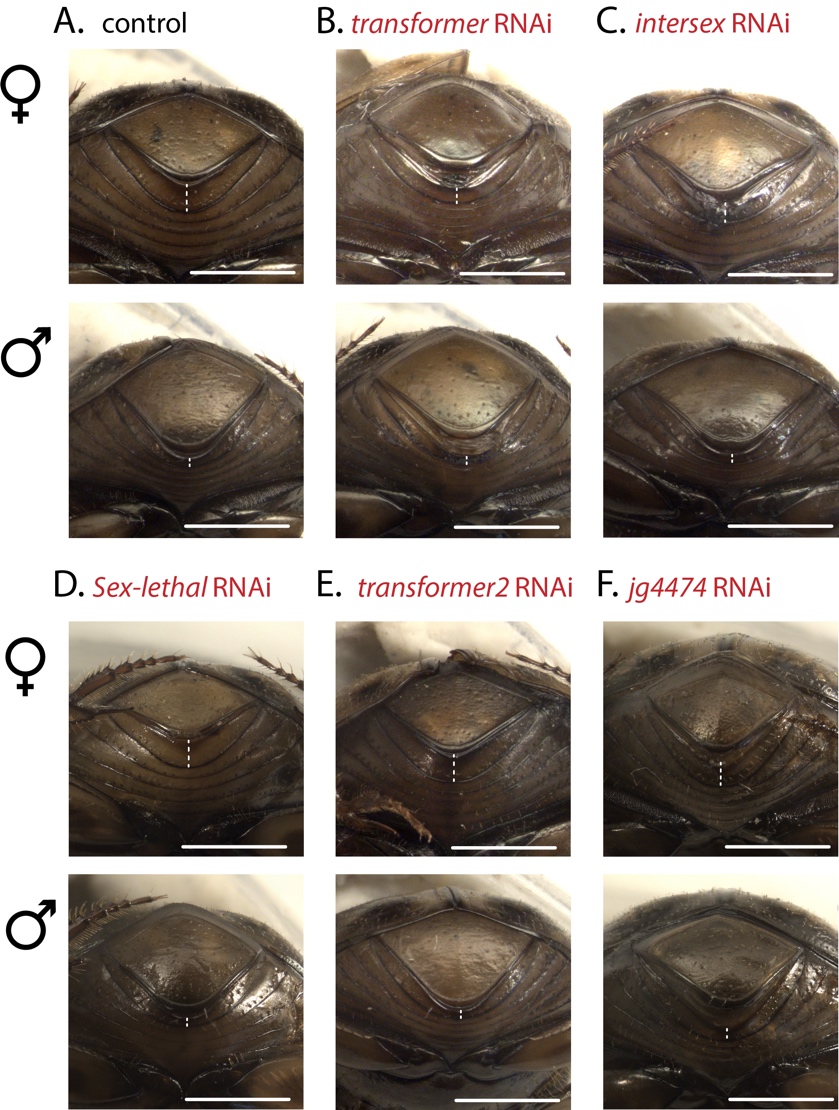

**Supplementary Figure S1. RNAi phenotypes of sexually dimorphic posterior ventral abdominal sclerites.** The pygidium and the ventral abdominal sclerite it closes against have a reliable sexually dimorphic configuration that is routinely used to sex adult animals across the Onthophagine clade. Specifically, in the female pygidium the distance between the edge of the pygidium and the sclerite it closes against is fairly consistent and widens slightly medially (white dotted line in A, top). In contrast, the male pygidium is slightly elongated, causing a conspicuous narrowing of that same distance medially (white dotted line in A, bottom). Representative animals obtained after control injections (A) and dsRNA injections targeting *Dg-tra* (B), *Dg-ix* (C), *Dg-Sxl* (D), *Dg-tra2* (E), and *Dg-jg4474* (F) are shown. *Dg-tra*^RNAi^ females (B, top) show a marginal narrowing of the sclerite compared to controls, indicating modest masculinization in this body region. *Dg-ix*^RNAi^ females (C, top) also show a marginal narrowing of the sclerite compared to controls, matching the degree of phenotypic change in other body regions. *Dg-tra*^RNAi^ males, *Dg-ix*^RNAi^ males, and both males and females of *Dg-Sxl*^RNAi^, *Dg-tra2*^RNAi^, and *Dg-jg4474*^RNAi^ show no change in phenotype, matching what was seen in other body regions. Scale bars = 1mm.

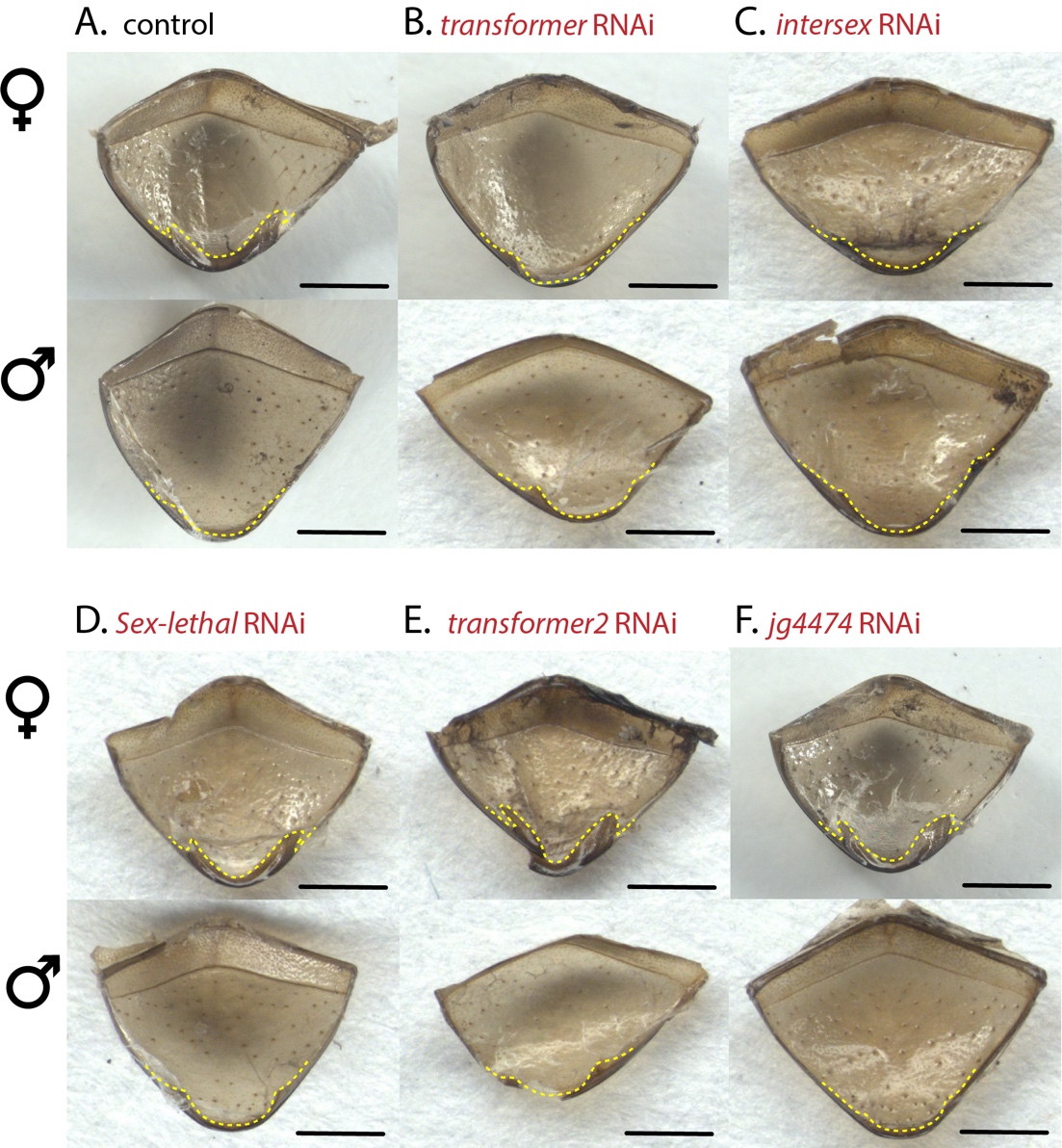

**Supplementary Figure S2. RNAi phenotypes of pygidium interior.** The cuticular grooves along the inside of the pygidium display a sexually dimorphic shape that can be used to sex adult animals across the Onthophagine clade; specifically, the female cuticular grooves have a more prominent bilaterally rounded shape (yellow dotted line, A, top) while the male cuticular grooves are smaller and protrude less from the edge of the pygidium (yellow dotted line, A, bottom). Representative animals obtained after control injections (A) and dsRNA injections targeting *Dg-tra* (B), *Dg-ix* (C), *Dg-Sxl* (D), *Dg-tra2* (E), and *Dg-jg4474* (F) are shown. *Dg-tra*^RNAi^ females (B, top) show masculizined narrow grooves compared to control females. *Dg-ix*^RNAi^ females (C, top) also show some narrowing of the grooves compared to controls, matching the degree of phenotypic change in other body regions. *Dg-tra*^RNAi^ males, *Dg-ix*^RNAi^ males, and both males and females of *Dg-Sxl*^RNAi^, *Dg-tra2*^RNAi^, and *Dg-jg4474*^RNAi^ show no change in phenotype, matching the phenotypes seen in other body regions. Scale bars = 1mm.

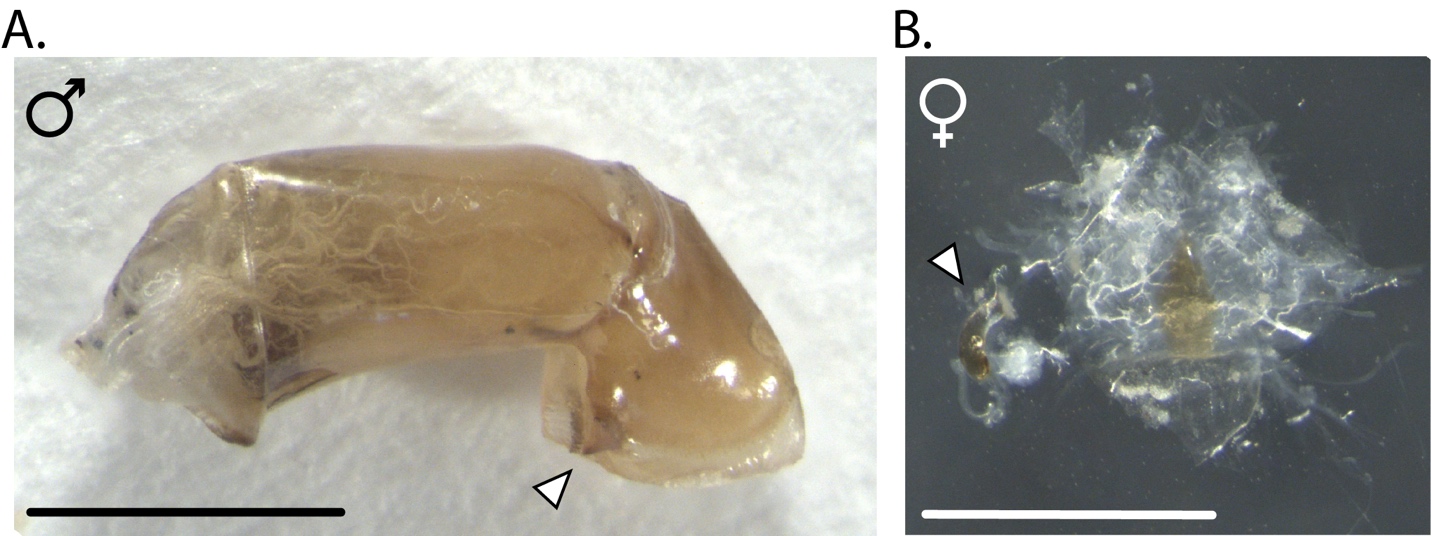

**Supplementary Figure S3. *D. gazella* adult internal genital structures. (A)** Dissected male aedeagus, positioned with proximal phallobase on left and distal parameres on right, hooks indicated with arrowhead. **(B)** Dissected female vaginal membrane and attached spermatheca indicated with arrowhead. Scale bars = 1mm.

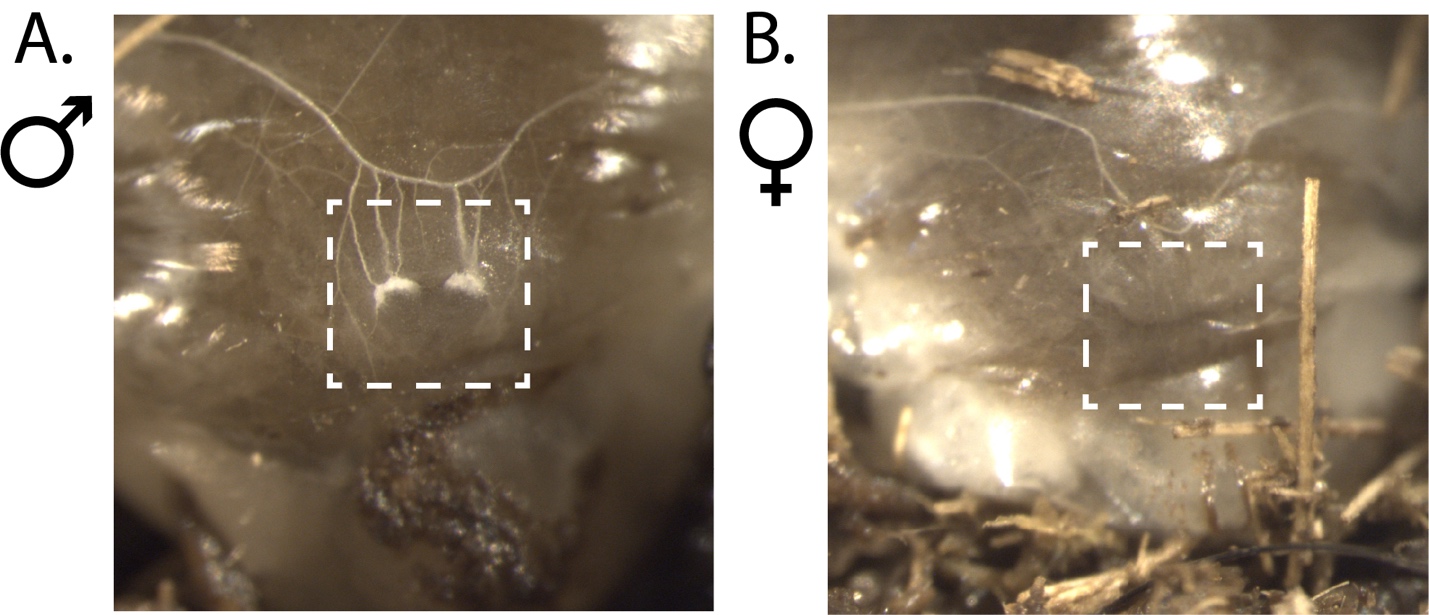

**Supplementary Figure S4. Visual sexing of *D. gazella* larvae.** Male Onthophagine larvae in the late second to third larval instar stages exhibit genital primordia visible through the ventrocaudal cuticle of the terminal abdominal segment (A, white box). These paired tissues are absent in female larvae (B).

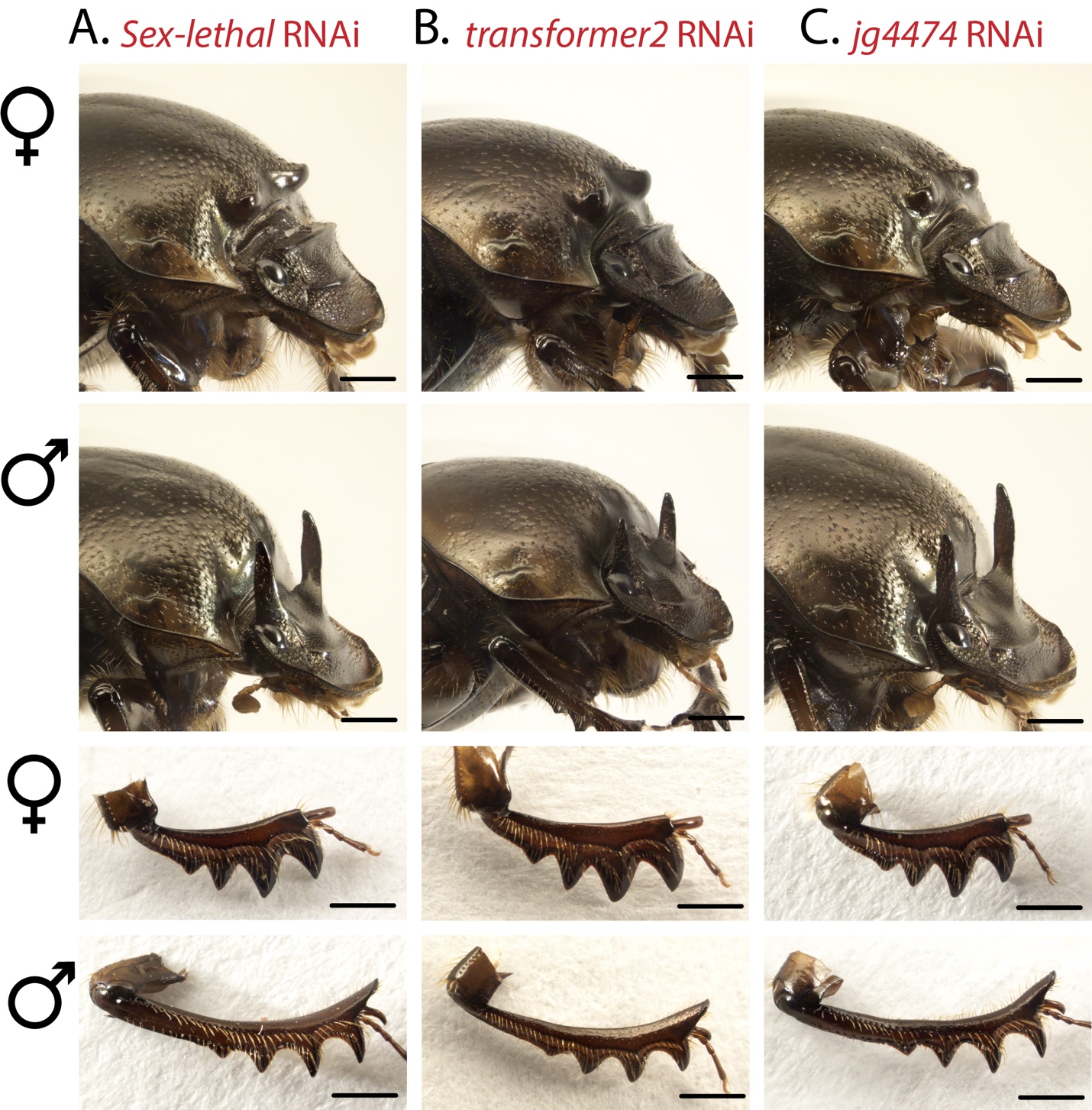

**Supplementary Figure S5. RNAi targeting *Sex-lethal, transformer2*, and *jg4474* has no obvious effects on D. gazella development.** Representative animals (top) and their dissected fore tibiae (bottom) obtained after *Dg-Sxl*^RNAi^ (A), *Dg-tra2*^RNAi^ (B), and *Dg-jg4474*^RNAi^ (a potential hermaphrodite homolog; C). These RNAi experiments did not affect adult male or female morphology. However, *Dg-tra2*^RNAi^ did result in high mortality (37 out of 41 dead). Note that the single surviving male *Dg-tra2*^RNAi^ individual shown in (B) is much smaller in overall body size than the rest of the size-matched males shown in all other figure panels; the length of the horns and fore tibiae can thus be accounted for by allometric scaling and does most likely not constitute an RNAi phenotype. Scale bars = 1mm.

**Supplementary Table S1.** BLAST Hit Sequences from *D. gazella* genome and RNAi target gene regions.

| **BLASTp Query** | **BLASTp Hit** | **Dmel query e-value** | **Dmel query bit score** | **Tcas query e-value** | **Tcas query bit score** | **Otau query e-value** | **Otau query bit score** | **Hit Gene Sequence (RNAi target region underlined)** |
| --- | --- | --- | --- | --- | --- | --- | --- | --- |
| Sex-lethal | jg9090.t1 | 2e-64 | 215 | 2e-97 | 286 | 2e-164 | 457 | ATGTTCACAAATAAGAAACTTGGACCTTTCCTTGGTTTTGGACAGACACGTGAATCCGCACCTATTAAAATGGACTGTCAGGAATCGAGTTCTAATAACGGTCATTTAACCAACCGTCCGACATCGTCAGGACAAAATGATGGATCCGGTGATGCCGATAACGTCGATAAAACCCGATTGATCGTTAATTACATCCCTCAGTTTACAACGGAGGAGGATTTAGCCCTCGTTTTTTCTCAAATCGGACCTATCGAGAGTATTCGAATTATGAAGAATTTACGTACCGGATATAGTTACGGATACGGTTTTGTTAAATATTTAAAGGCTGAAGATGCCGCTAAAGCCATCGAAGCACTTAGCGGATTAAATTATAGAAATAAACGCCTAAAAGTGTCATATTCGAGACCTCCCGGTCAAACGATGAAAGATTCTAATTTGTACATCTCGAACTTACCCAAAGACGTTACCGAAAAGGATTTAGACCGAATGTTTGGTGTTTACGGAGAGATTATCCAGCGGAATGTGTTAAAGGATAAGATTACTGGGTTGCCGAGAGGTGTCGCTTTTGTACGGTTTTCGAAGAGAGAAGAGGCTCAAGCGGCTATAGCCAGTTTAGACGGTAAACAACTTGAGAAATCGATGTTTCCTTTAAGCGTGAGAGTCGCGGACGATCACAGCAAACAAAAAGCTCAGTTTTTAGAACACCAATCGTGCTTTTTGGGCAGAGGTTTTGGCGGCTTCAGGGGTCCGTTTCCGCGCAATGGAACGTTTCCGGTTCAAAGGAGCCCTTCTAATTTCGTGAGCAGTCGATTTTTGGGAGCGGGCGACAGCCACTACCTCAGCTCGATGTTTTGGTAG |
| Transformer* | jg4056.t1 | - | - | - | - | 1e-86 | 269 | ATGGAACCAAATAAACGACCTCTAAAAAGATTACCGCAAGTAACTTTATGGGAACTTCAAAGACACACTATAACCGTAAAAAATCCTTTGGCTTGGGATAAAACCGGCTCGGATAAACCAATACAACGCGATATAGTGGATTTAAGTGAATTATGTTTACACCGGGCAGAGGTTTATAGTAAGAAAAGGATTAATAGATGGTTACTAAAGGGTTTTTGGTGGGATTCAAGAGGAAACCAACAATGTTGTGAAGGTTCGACACCGCTTTTCGATAGAGGTGATATCCTGGTGAATCAAGATGGGATGGAAACGCGTACAATGAGGATTAAACGTACAGGACGATCGCGTTCCTCTTCCAGTTCATCAAGTTCAGATTCATCACGATCCAGACGGAGTAGCAGCCGTACAAAATGGAAAACCGAGGCTAAAATCAACATTCCAAAACCACTTAGGTTCACTTGCGAGAGAAAACGTTCACCAGTTAATCAAAAAGATGACTCACATAGATTGAGGCGTAATTCGGATCGAAAATCGGTTTCTCCAAGAAGAAGACCAAGTATTTCACCACATAGAAGAATGACTAAGCGGAAATCGAGTAACTCTCCGAGTAAATCTCCGTATCGAGGGGATAAAAGGATTTCTAGAGCTGAAAGATCGCCCGTTAGGAGGAATAGAAGTCCTTTAAGGGGTGGGAAATCCCCTGAAAGAGCGGGAACATCTCATATGAAAGATGAAAAGTCGCGTAGATCGGAAAAGTCGCCATTAAGGTCGAGGGACAGAAGAAGGAGTCGATCTAGAAGTAGATTGAGGAGAAATACTCCTCCGGGGCGGAGTAGATCGCCTCGGAGAACTAGTAGTACGAGACATAGAGAGACCGGAACTTGGAATAGGTCTCCTAATAGGCAAAGAAGAGATTATTCACCTCATAAAAGTGGTGAACATATTCCTCCTATTGATTATGAACATTTAGCACATGTTATCACCCAGATGTATCCAAAACAACCATTTTTTACACCACCTATAGTACACCCACCTATGGCAGAGTTTTATCCGCCACCATATGGGATTCCTCCGGGGCCTCAATTGCCGCCAAGACCGAGATTTCCACCAAGACCACCTATAATCTACAATAATAGAACGATAAAAATCTTTAGACCCAACGAAACGACATCCACATCGAATACAACTACAACAACTGTTACTACTACAACCCCGATAACAAATACGCGACAAAATCCCGGTTCTTCAACGGTAACTAGTAACGAACCGGATTCGGAAGTTAGAAATGATGTAGATAAATAA |
| Transformer2 | jg25106.t2 | 5e-75 | 228 | 5e-75 | 228 | 5e−324 | 519 | ATGAGCGACAGAGAGGCTACTAGGTCTCCTGGACACAAACCTATTAACGATCGTGATCATTCGCCTCACGGAAGGCGGGAAGAATCACCTAGGAATCACAAGACAAAAAGCTATTCTAGAAGTCCAAGTCGTGGGAGAAAAATGTCTTACAGAGACGATCGGGACAGAGAACGTCGCAGGGATGGAGACCGCGATGGTCGTGGTGACGGTGGACGACGTTATAGGTCCCGCAGTCGATCACGCTCAGACAGACGTTACAAGTCCTCTCGATACTCCCGCAGCCGGTCCCGCTCAGGTGGTTACGGCCGACGATCCCACTCACATAGCCCTATGTCTAGCCGTAGGCGCCACATGGGTACCCGCGACAACCCGAAGCCGTCCCGATGCCTGGGCGTTTTCGGTCTCAGCGTCTACACAACCGAAGACGAGCTTTACCACATTTTCTCCAAATACGGACCCGTTGAGCGCGTGCAAGTTGTGATCGATGCTAAGACTGGTAGGTCGAGGGGATTCTCCTTCGTTTACTTTGAAAATACCGAGGACGCTAAAATCGCCAAGGAGCAATGTACGGGGATGAAAATCAACGGGAAGAGTATCAGGGTGGATTATTCGATTACCGAACGTGCTCACACTCCAACGCCCGGTATTTATATGGGCAAACCAACATACATCAGCGATAAATGGGATAGGCGCAGAGATAGGGATAGAGATGATTATTATGGAGGTGGAGGATACAGAGGTTCCAAATACAGTCACAGAAGGTCTCCGTCACCATATTACAGGCGACGACGCTACGATCGTTCCAGATCAAGATCGTATTCTCCTCGGTTAAGAACACGTGGTATTGGATGA |
| Intersex | jg1008.t1 | 6e-40 | 133 | 2e-76 | 225 | 1e-116 | 327 | ATGAACGTAGTTGGAATGGGAATGCCCCCTCAACAAGCCCCCCAACCGGCTCAACAACCTCAACAGCAACAGCCGCCGCAATCTCCGATGGATAACATATCAAAAATTAAAACATTAATAGGACCTTTACGTGAAGCACTGAACAATACCATCAAAACTGCGGCACAAACTTTAAATCAAAATAGCCAGGTGGATATAGGCTCACAAAGGGGTGTTGATCAACAAATTCCACGGTTTGATAAAAATTTGGAAGAGTTTTATTCACTTTGCGATCAAATAGAACAAAACTTGAAAACCTCAATTAAATGTTTGAGTCAACATGAAAGTGCTCAAAGATATTTGAATCTCCTGGTAGCCCCTACCAAAAGTGAAAGTATTGGAATTCCTGATAATACTCTTTCTTATCCGCAATATTTAGCTACGGTTAATGCCCAAATAGCGTTTACTAAGGAAATACATGATACACTGGTAGCAGCAGCTCAAAACATATCCCCATCTGAATGA |
| Hermaphrodite | jg4474.t1 | 5e-04 | 42.4 | 6e-23 | 101 | - | - | ATGAGTATCAAAATAGAAAATGAATTGAGGCCCCAAACACCTCAAGAATCCGAGACGTCTTTTTCTTCAAATTCAAGTTTTATACCGAATTTACCCCAAGAAAATCAAGATAATTGCGAGATCAAAATAGAAGAACAAGAAGTCTTCGAAGGGTTTGAAACCCCAACTTTGAGCTTTGAGGAAACTTCTATCGTCAAATATCCATTTATACAAAGTTCTCTTCATACTACAAGCTTGGGTGAGGCTTGTAATGAAAGTACGGGTATTTACGTTTGTTCATTTTGCAATGTTGAGATATCGGAAAAGAAGTATTATGGTAGGCACATAAAATGGTGTGCTAGAAATTATTTAATTATGAGTAAACAAGAGGAGTTCACCACAAAAGTAAAACGGAAATTCAACTGCGACAAATGCAGCTACTCAGCAAAAAATAATTACAACTTACAACGTCACGCTTTAGTTCATACCGATCGTAAACAATTTAGATGCGTTATTTGTAATAAATCCTTCCGACGAAACAGTCCTTTACGCGATCACATCAAAACCCACAACACCAATCGAGAAAAAAACGTTTTATGCCCTTATTGTAAAAATACCTACCATCGTAAAAGCGCTTTAACGCGTCACATCCAAGTAAAACATTCCGATCTTATTAAATTCAAATGCGATTTATGCGAATACACAACCATCGATGAAATCCAACTTAAAAAACATATATCTTCCAAACATACCGATTTTAATAAAAAATACGAATGCGATCATTGCGATTATTTTACCGAAAATAAAGGCCACATGAAAACCCACCTTAGAATACATACAAACTTAAAACCCTTTAAGTGTAATTTTAAAATATGTAAATGTTTAAAAAACATAGAAAAATTATGCGAATGCGAATGCAAACCATGTAATTTTGAAACCGCACACGAAGTTAATTTAAAAAAACACGTGCTTCAAAAGCACAGTAACGAAACGAAAAAATATAAGTGTTCCGATTGTTGTTATTCCACACACATAAAAGCGGATTTTACGACTCATTTAAGAATACATACGAATGAAAAACCTTTTAGTTGCGGATATCCTAATTGTGAGTATAAATCGACTCGAAGGGAACATTTAAAAATACATATTCGACATAAACACACTAAGGAGAAACCGTTTAGGTGTGAAATGTGCGATTATTCGTGTGTTAGTAGTTGTAATTTAAAGAGACATGTTATCAAGCATAATAATTAA |
| Hermaphrodite | jg1708.t1 | 9e-06 | 48.1 | 4e-19 | 91.3 | - | - | ATGTCTGCACAGTTTGGACAAACACAGATCAAACTCGAGGGGACAGACTCCGCGGAGAATGTCACCGAGATACAATCGTACCTCGAGGGGTTTCAAAAGGAAATCGAAGGCACCGATAATAATACAACTCAACAAGTTAAAGAAATAGATGATGACAATGATGGTGATGGAGACGAAGGAACTTACTTTGTGGACCAAGCCGGTCATTATTACTACCAGGCAAAGGGAGAAACTCAGCCTGTTATGACTATGGTTTCGGGGATCAATGATTCAGAAGGTGGTGATGGTGAAGAGTTTATTATCAATCATGAAAATGATGAAGATGCAGAAGAAGATGGCGAAGAGGGATTGCCTGGCGATGGAGAAAACAACCAGATTCTTATAAATAATGGAAACGCATACCAAAGAATAACAGTCGTTCCTGCAGACACAAGCTCCAATGAGTTGAGCTATGTGCTTATCGTTCAACAACCAGATGATAAAGAAGGGGAACAAACTGATGGAGATCAAGATATGGCCGTTTACGATTTCGATGAAAACGAAGAAGGAAACGTAGTCGATTCGGAGGCAGAAGATGACAAGTCAAAAATAGTAAAGCTCCTCCCAAGACGATCTCAAGTTGTTTCTCAGCCGTACATGTGTAACTATTGCAATTACACCAGTCCGAAAAGATACTTGCTCTCGCGTCATATGAAATCACATTCGGAAGAACGGCCTCATAAGTGTAGCGTTTGCGAGCGCGGTTTCAAAACGGTCGCTTCCCTCCAAAACCACGTTAACACGCATACGGGTACTAAACCGCATCAATGTAAATATTGCGACGCCGCATTTACTACTTCCGGCGAACTCGTCAGACACGTCCGCTATCGTCACACGCACGAAAAACCGCACAAGTGTAATGAATGCGATTATGCCAGCGTTGAACTATCGAAATTGAAAAGACACATTAGGTGTCATACTGGTGAAAGACCGTATCAGTGTCCTCATTGCACTTATGCAAGTCCTGATACTTTTAAATTAAAGCGACATCTTCGTATTCATACCGGTGAAAGACCGTATGAATGTGATATTTGTCAGGCGAGATTTACCCAGTCGAATAGTTTGAAGGCGCATAAGTTAATTCATAATGTTGGCGATAAACCGGTATTTCAATGCGAGCTTTGTCCGACAACTTGCGGTAGAAAAACCGATTTGAGAATACACGTTCAAAAATTACACACTTCGGACAAACCGCTTAAATGTAAAAGATGCGGGAAATCGTTTCCCGATCGGTACAGCTATAAACTCCATAATAAATCCCACGAAGGGGAAAAATGTTTTAAATGCGATTTATGTCCGTACGCTTCGATTTCCGCCAGACACCTCGAATCGCATATGTTGATCCATACCGACCAAAAACCGTTCCAATGCGAACATTGCGATCAATCGTTTAGACAAAAGCAGTTACTTAAACGCCACATTAACCTTTATCATAACCCGCTTTACGTACCGCCGACGCCTAAAGAAAAAACCCACGAATGTCCCGAGTGTCACAGACCGTTTAGACACAAAGGAAACTTAATCCGACACATGGCGGTTCACGATCCCGACTCTAGCATACAAGAGAAGCAATTAGCTTTGAAACTCGGCAGACAGAAGAAGATACAAATGATCGACGGTCAACAAGTCGAAGTTATGCCGAGTATGGGCTCAGATGATGAGGAGTCTGATATGATGGCTGTTGAAGGGTCGGATGGACAACAATATGTTGTTTTGGAGGTTATTCAACTTGCTGATGGTGAGGAGCAGGCTATGGTTGTTGACGGACAAAGTGATATGCTTGGAGAAGGAATTCTACAAGATGATTTACATGATGAAGAAGTTATCAAAGCTCTTCAAAATGCTGCTAGAAGAGGAGTAACCAAAGTTGAACAAGAAGACGCTGAAGAAGAGGAAGAAGAAGATGATAAGAAAATGGACCATGATATGGAAACTTGTTTTGGTTTTGATGAGGAAGAAGAAGAAGATGAAGATATGAGACATGGAAAAGAAACAATAACCCTATTAGGAATGGACTGA |

* *Trypoxylus dichotomus* query also used for Transformer; e-value = 0.002, bit score = 40.4

**Supplementary Table S2.** Primer sequences and usage

| **Primer** | **Sequence (from 5’ to 3’)** | **Use** |
| --- | --- | --- |
| *Sxl-forward* | TAATACGACTCACTATAGATGTTCACAA | dsRNA preparation |
| *Sxl-reverse* | TAATACGACTCACTATAGATTTGAGAAA | dsRNA preparation |
| *tra-forward* | TAATACGACTCACTATAGATGGAACCAA | dsRNA preparation |
| *tra-reverse* | TAATACGACTCACTATAGTGATTCACCA | dsRNA preparation |
| *tra2-forward* | TAATACGACTCACTATAGATGAGCGACA | dsRNA preparation |
| *tra2-reverse* | TAATACGACTCACTATAGCCTGAGCGGG | dsRNA preparation |
| *ix-forward* | TAATACGACTCACTATAGATGAACGTAG | dsRNA preparation |
| *ix-reverse* | TAATACGACTCACTATAGGTTTTCAAGT | dsRNA preparation |
| *jg4474-forward* | TAATACGACTCACTATAGGAGGCCCCAA | dsRNA preparation |
| *jg4474-reverse* | TAATACGACTCACTATAG CGTAAATACC | dsRNA preparation |
| *jg1708-forward* | TAATACGACTCACTATAGATGTCTGCAC | dsRNA preparation |
| *jg1708-reverse* | TAATACGACTCACTATAGACAGGCTGAG | dsRNA preparation |
| *dsx-forward* | ATGTCTGATCAGCAGGATTAC | RT-PCR |
| *dsx-reverse* | TCACGCTCTACTTCTAGGCC | RT-PCR |

**Supplementary Table S3.** RNAi treatment sample numbers and penetrance data.

| **RNAi Target** | **# Injected Larvae** | **# Dead** | **# Surviving Adults** | **% Mortality** | **# Adult Males** | **# Adult Females** | **Penetrance of Phenotype (females only)** |
| --- | --- | --- | --- | --- | --- | --- | --- |
| *Dg-ix* | 42 | 16 | 26 | 38% | 14 | 12 | 100% |
| *Dg-tra* | 35 | 14 | 21 | 40% | 13 | 8 | 90% |
| *Dg-tra2* | 41 | 38 | 3 | 93% | 1 | 2 | - |
| *Dg-sxl* | 21 | 5 | 16 | 23.8% | 9 | 7 | - |
| *Dg-jg1708* | 26 | 26 | 0 | 100% | 0 | 0 | - |
| *Dg-jg4474* | 26 | 4 | 22 | 15.4% | 13 | 9 | - |
| controls | 33 | 13 | 20 | 39.4% | 10 | 10 | - |

**Supplementary Table S4.** *D. gazella doublesex* isoform sequences

| Male isoform | ATGTCTGATCAGCAGGATTACNNNNNNNNNGGCAAGGTCTGAATGCGTCTAGCACATCGAGTAGCCCTAGAACGCCACCGAATTGCGCCCGTTGCAGGAATCACCGGNGTAAAAGTGCCCCTAAAAGGCCACAAGCGGTACTGTAAGTACAGACATTGCAAGTGCGAGAAATGCCGGCTTACCTCTGAAAGACAGAGGGTGATGGCCATGCAGACCGCCCTGCGAAGGGCTCAAGCTCAAGACGAAGCTATGTTACGGCAAGGAGTGATTCCACAACCGAAAAGTCCCGTACCTCTTCATGGCACCGATCGAACATACGAGTGTGAATCCCCGGGATCGTCTTCAACGCCGACCTTTCCAGAAGTCATCCGGAAAACTCCGATGGAACCCATTAGGGATACAATGGTGAATAACAATGTTGGACAAAAAGATTTAATACAAGAAAGTTTACAACTGCTCGAAAGGTTCCGATATTCATGGGAAATGATGCCATTGATATATGCAATAGTTAAAGACACTCCAGACCTTGAAGAAGCGTCAAAACGTATAGATGAAGGTAAAGATGCTGAGCAGCTATTGGATTTTTTCAATAAAATAAAGGACAGGTTTCACCTATCCTGGAAAATGATCTCACTTATCCACGTCATCCTGAAGAACGCAAAGGATGATCAGGAGAAAGCCTTTAGACAAATAGATGAAGCATTCCTGGAGGTGCAGACCTTAGCAAAATACTACCCAACACCAGTGAATCCAATCACTGACTTATGGAGGTGGTATTCCCCTACAGCACTCTACCCAGCGATGTACTACCAGCTGCCCAATACTCTTCTTGGAAGTATTCCTCCAAGTCCACCTTTACACTCGCCAGTACCACCCCCACGGCCTAGAAGTAGAGCGTGA |
| --- | --- |
| Female isoform 1 | ATGTCTGATCAGCAGGATTACNNNNNNGGGAAAGGTCTGAATGCGGTCTAGCACATCGAGTAGCCCTAGAACGCCACCGAATTGCGCCCGTTGCAGGAATCACCGNNGTTAAAAGTGCCCCTAAAAGGCCACAAGCGGTACTGTAAGTACAGACATTGCAAGTGCGAGAAATGCCGGCTTACCTCTGAAAGACAGAGGGTGATGGCCATGCAGACCGCTCTGCGAAGGGCTCAAGCTCAAGATGAAGCTATGTTACGGCAAGGAGTGATTCCACAACCGAAAAGTCCCGTACCTCTTCATGGCACCGATCGAACATACGAGTGTGAATCCCCGGGATCGTCTTCAACGCCGACCTTTCCAGAAGTCATCCGGAAAACTCCGATGGAACCCATTAGGGATACAATGGTGAACAACAATGTTGGACAAAAAGATTTAATACAAGAAAGTTTACAACTGCTCGAAAGGTTCCGATATTCATGGGAAATGATGCCATTGATATATGCAATAGTTAAAGACACTCCAGACCTTGAAGAAGCGTCAAAACGTATAGATGAAgggcaaagagcagtgaaggaatactcaataatcaacaacctcaacatgtacgacggtggagaacttcgttatcctactcgatgaaggatgcaagaacgacgaccatccgggaacttcaaatccgtcgacggcattaacctccaggtgaaatgcaacttacacgttacataggatcatctcatttaccgtttagtaaaaatttttagcggttaacgtttcacacttggcattaaatgcgataaccgaaaacgcctcagcgagtttatgaaataaatacaataagaaacgttgaactgtatagtattataattattaagtattaagcttgtattgagtcagtattcatattttctgttgacgaagcctcaatttataacagtattgtgcattttacgacatttgtatatttctctacacaactaccacagaaaaaaaaatGGTAAAGATGCTGAGCAGCTATTGGATTTTTTCAATAAAATAAAGGACAGGTTTCACCTATCCTGGAAAATGATCTCACTTATCCACGTCATCCTGAAGAACGCAAAGGATGATCAGGAGAAAGCCTTTAGACAAATAGATGAAGCATTCCTGGAGGTGCAGACCTTAGCAAAATACTACCCAACACCAGTGAATCCAATCACTGACTTATGGAGGTGGTATTCCCCTACAGCACTCTACCCAGCGATGTACTACCAGCTGCCCAATACTCTTCTTGGAAGTATTCCTCCAAGTCCACCTTTACACTCGCCAGTACCACCCCCACGGCCTAGAAGTAGAGCGTGA |
| Note: Regions targeted by primer sequences in Table S3 underlined in black. Sequence resolved by Sanger sequencing in uppercase. Sequence resolved by Illumina sequencing in lowercase. | |
